# STING activation depends on ACBD3 and other phosphatidylinositol 4-phosphate-regulating proteins

**DOI:** 10.1101/2022.10.17.512580

**Authors:** Rutger D. Luteijn, Sypke R. van Terwisga, Jill E. Ver Eecke, Liberty Onia, Shivam A. Zaver, Joshua J Woodward, David H. Raulet, Frank J.M. van Kuppeveld

**Author notes:** equal contribution (David H. Raulet, Frank J.M. van Kuppeveld).

## Abstract

STING induces transcription of pro-inflammatory genes upon activation at the Golgi apparatus. Many of the regulators involved in STING activation are unknown. We found that ACBD3 and other phosphatidylinositol 4-phosohate (PI4P) regulating proteins play a critical role in STING activation. We show that proper STING localization and activation at the Golgi depended on ACBD3 and PI4KB expression. Furthermore, depleting PI4P by inactivating PI4KB or overexpressing Sac1 diminished STING activation. STING signalling was also regulated by the lipid-shuttling protein OSBP, which removes PI4P from the Golgi. OSBP inhibition by the FDA-approved antifungal itraconazole and other OSBP inhibitors greatly enhanced STING activation by increasing the levels of STING-activating phospholipids. Itraconazole-enhanced STING activation resulted in a hundred to thousand-fold increased expression of interferon-beta and other cytokines. In conclusion, the phospholipid PI4P is critical for STING activation and manipulating PI4P levels is a promising therapeutic strategy to alter the STING immune response.

## Introduction

Cytosolic DNA is a key danger signal that can be detected by various cytosolic DNA-sensing pathways, most notably the cGAS/STING pathway. This innate immune pathway senses cytosolic DNA originating from viruses and bacteria^1^ as well as cyclic dinucleotides (CDN) produced by certain bacteria^2–4^. The STING pathway is also activated by cytosolic self-DNA, which accumulates in cells in certain autoinflammatory disorders^5,6^, and in cells subjected to DNA damage, as occurs in premalignant and tumour cells^7,8^. In addition, the cGAS/STING pathway plays a role in the immune response to certain RNA viruses, such as dengue virus^9^, influenza virus^10^, and coronaviruses^11^. RNA viruses may trigger the cGAS/STING pathway by stimulating the accumulation of host DNA in the cytosol of infected cells^12^. Moreover, STING can be activated by virus-induced lipid membrane remodelling events^10^. The critical role of STING in the immune response to virus infection is underlined by the observation that numerous viruses, including, herpes virus, vaccinia virus, dengue virus and SARS-coronavirus^11^, counteract STING activation, thereby evading the host immune response.

STING regulation is a complex process that starts with binding of ER-localized STING to its ligands, most notably cyclic dinucleotides (CDNs). The mammalian CDN 2’3’-cGAMP is produced endogenously by the enzyme cGAS upon detection of cytosolic DNA and binds STING with nanomolar affinity. In addition, 2’3’-cGAMP can be imported from the extracellular environment or neighbouring cells to activate STING^13^. Similarly, synthetic phosphodiesterase-resistant CDNs, like 2’3’-RR CDA (RR CDA) used in cancer immunotherapy, are transported into the cell and bind ER-localized STING with high affinity (nanomolar range)^14^. Upon CDN binding, STING dimerizes and translocates to the Golgi compartment by a poorly understood process. At the *trans*-Golgi network (TGN), STING oligomers are phosphorylated by TBK1, and STING subsequently activates the transcription factors IRF3 and NF-κB. To prevent sustained immune activation, activated STING is degraded in the endo-lysosomal compartment^13^.

Anomalies at any of these steps can lead to aberrant STING activation, resulting in auto inflammatory conditions^15^, or diminished STING signalling and immune escape, as observed in certain tumours and virus-infected cells^16,17^. Furthermore, STING activity can be redirected to generate a tumour- or virus-promoting environment^18,19^. Many of the factors orchestrating the quality and intensity of the STING response remain unknown.

To find factors regulating STING activity, we recently performed a genome-wide CRISPR interference (CRISPRi) screen^20^. Using this method, we successfully identified transporters that import STING agonists from the extracellular environment. In addition, we identified many host factors that may drive or dampen STING activation. One of the top hits in this screen that was necessary for strong activation of the STING pathway was the gene encoding ACBD3. This Golgi-resident protein is a multifunctional protein that regulates the distribution of the phospholipid phosphatidylinositol 4-phosphate (PI4P) to the Golgi by recruiting the PI4P kinase PI4KB^21^, and has not previously been implicated in STING activation. Here, we show that ACBD3 regulates STING activation by affecting STING mobilization to the Golgi. We also implicate in STING signalling other components involved in PI4P biology, including Sac1, PI4KB and the PI4P-cholesterol exchanger OSBP, underscoring the importance of PI4P in STING signalling. The phospholipid PI4P plays a major role in recruiting several Golgi-resident proteins to the Golgi and the function of those proteins^22^, and may similarly be involved in STING recruitment or retention in the Golgi and STING signalling in the Golgi. The role of the PI4P pathway in STING signalling is especially interesting, as it provides novel therapeutic potential to modify the immune response by STING, including by targeting the pathway with FDA and EMA-approved drugs.

## Results

To confirm the role of ACBD3 in STING activation, we depleted the expression of *ACBD3* in THP-1 monocytes using CRISPRi gRNAs (Figure S1A). To measure STING activation in these cells, we expressed an ISRE-IFNb-tdTomato reporter, which robustly expresses tdTomato upon STING activation^20^. Indeed, control cells stimulated with the highly potent STING agonist 2’3’-RR CDA expressed the fluorescent tdTomato reporter, but not upon depletion of IRF3, a transcription factor critical for the expression of the reporter. In ACBD3 depleted cells, reporter activation by RR-CDA was also highly diminished, whereas STING-independent reporter activation by human interferon-beta was not affected (Figure 1A and 1B). Restoration of ACBD3 expression rescued reporter activation in ACBD3-depleted cells (Figure 1C).

**Figure 1.**
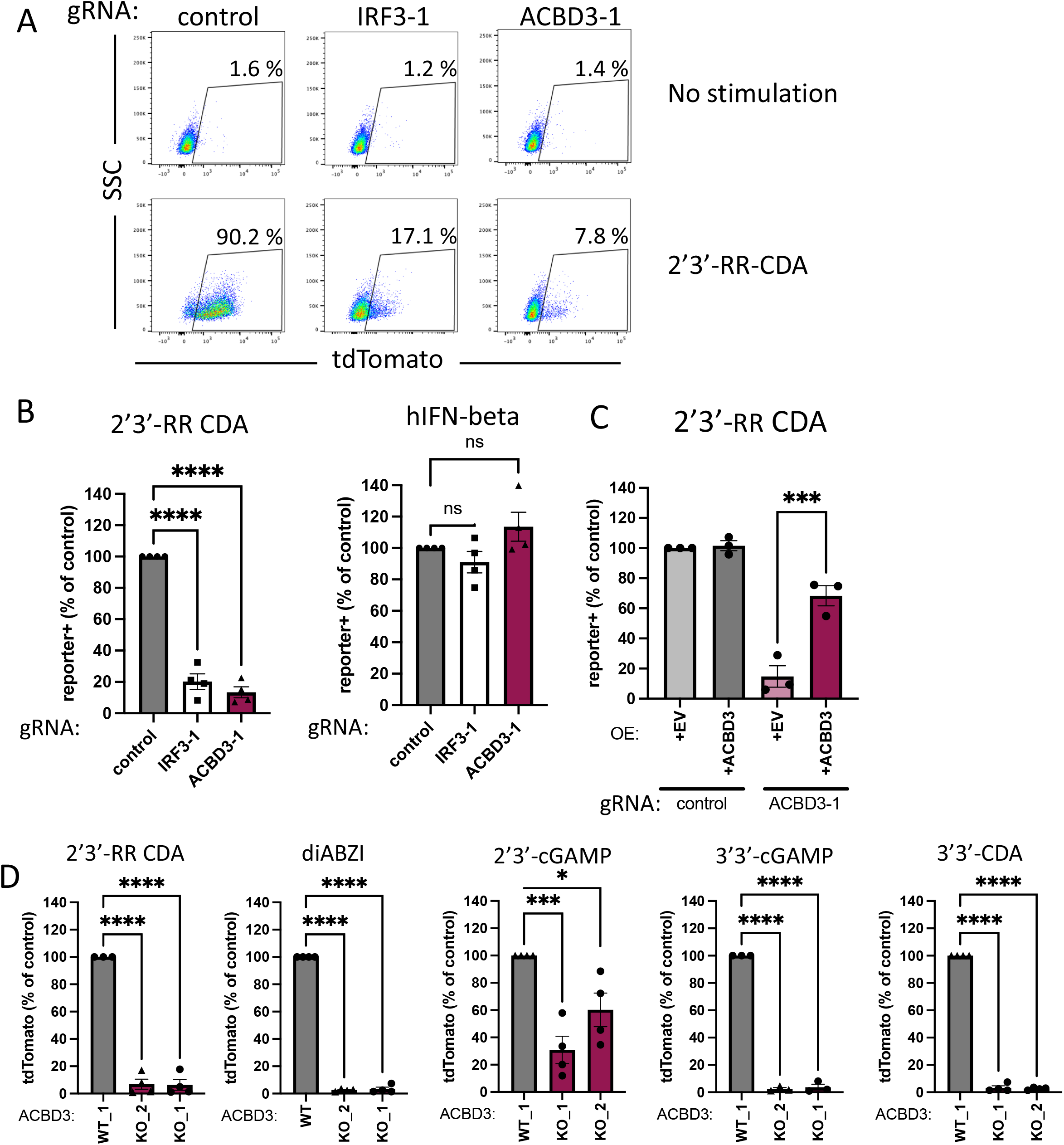
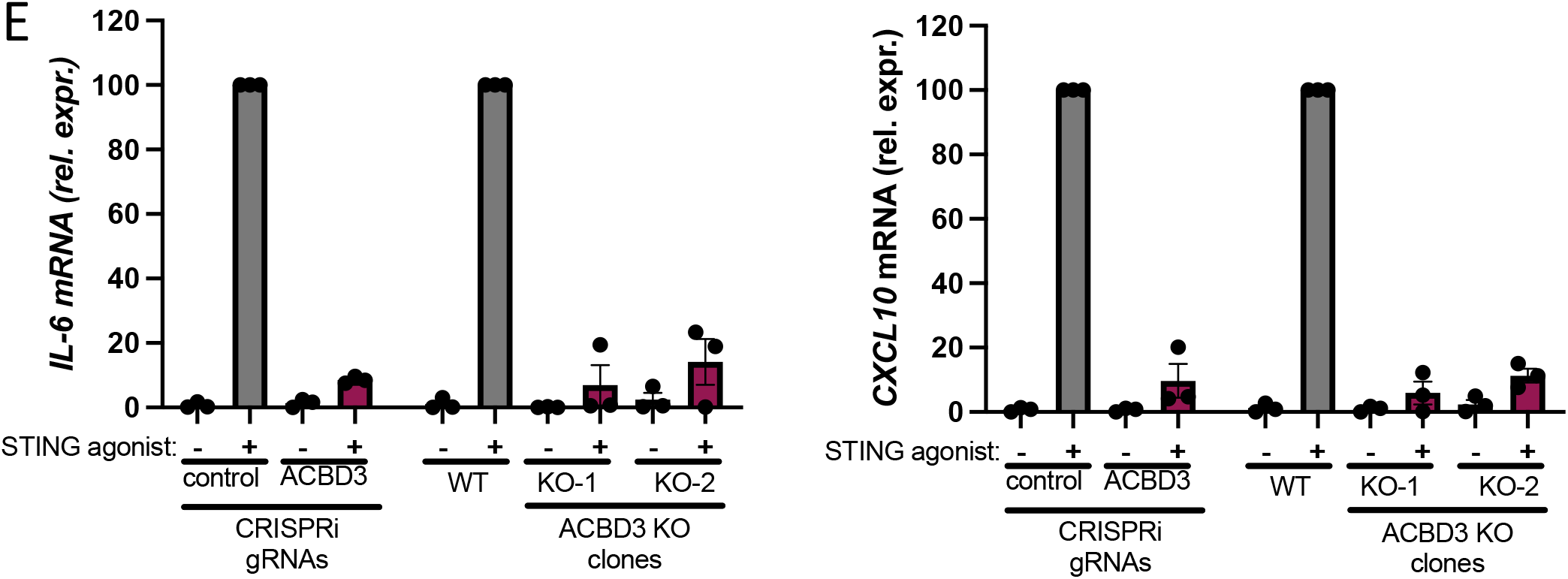
ACBD3 expression is necessary for tdTomato reporter activation and cytokine production induced by STING agonists a. dCas9–KRAB-expressing THP-1 cells transduced with non-targeting gRNA (control), IRF-3-targeting gRNA (IRF3-1) or ACBD3-targeting gRNA (ACBD3-1) were exposed to 2′3′-RR CDA (1.67 μg/ml). Twenty hours later, tdTomato expression was analyzed by flow cytometry. Representative dot plots of three independent experiments are shown. b. THP-1 cells expressing the indicated CRISPRi gRNAs or non-targeting gRNA (control), were stimulated with 2′3′-RR CDA (1.67 μg/ml) or human interferon beta (hIFNb, 100ng/ml) After 18–22 h, tdTomato expression was quantified as in a. c. Control THP-1 cells and THP-1 cells expressing ACBD3-1 CRISPRi gRNA transduced with ACBD3 or empty vector (EV) were stimulated with 2’3’-RR CDA and 20-24h later analysed as in a. d. THP-1 control clone (WT1) or two THP-1 clones lacking ACBD3 were stimulated with the indicated STING agonists and analyzed 20-24h later as in a. e. *IL-6* or *CXCL10* mRNA levels in THP-1 cells expressing control or ACBD3 CRISPRi gRNAs, or THP-1 WT or ACBD3 KO clones stimulated with 2’3’-RR CDA (RR-CDA) for 5h. b-e. Mean ± SEM of *n* = 3 biological replicates are shown.

As further confirmation, we generated ACBD3 knockout cells in THP-1 cells using the conventional CRISPR-Cas9 system (Figure S1B). Two distinct knockout clones lacking *ACBD3* expression showed highly reduced reporter activation to a variety of STING agonists, including 2’3’-cGAMP, 2’3’-RR-CDA, the bacterial CDNs 3’3’-cGAMP and 3’3’-CDA, and the non-cyclic dinucleotide STING agonist diABZI (Figure 1D). Similarly, *ACBD3*-knockout 293T cells (transduced to express eGFP-mouse STING, eGFP-mSTING) had reduced reporter activation upon STING activation (Figure S2). STING activation leads to downstream transcription of inflammatory genes, including *IL-6* and *CXCL10*. Expression of both genes was highly reduced in *ACBD3* knockdown and knockout THP-1 cells (Figure 1E).

To further investigate the role of ACBD3 in STING activation, we determined the effects of ACBD3 depletion on the different processes involved in STING activation. The uptake of 2’3’-cGAMP or 3’3’-CDA from the extracellular environment was not affected in cells lacking ACBD3, in contrast to cells lacking SLC19A1, one of the CDN transporters (Figure 2A). A second critical step in immune activation by STING is STING phosphorylation at S366, which is required for downstream activation of IRF3. In the absence of ACBD3, STING phosphorylation upon stimulation with 2’3-RR CDA was highly reduced (Figure 2B), whereas IRF3 depletion had no effect on STING phosphorylation. As expected, depletion of the SLC19A1 transporter also diminished STING phosphorylation.

**Figure 2.**
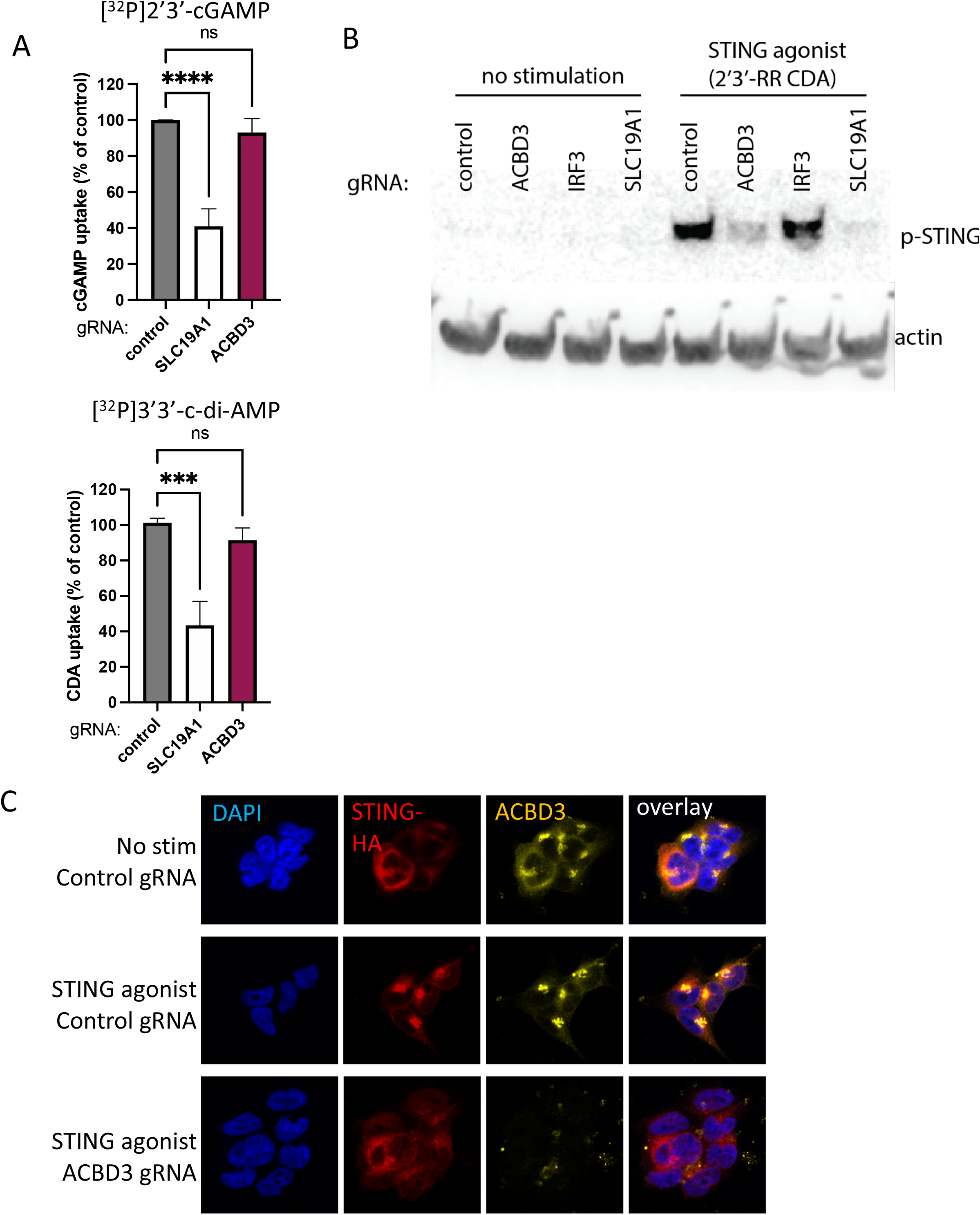
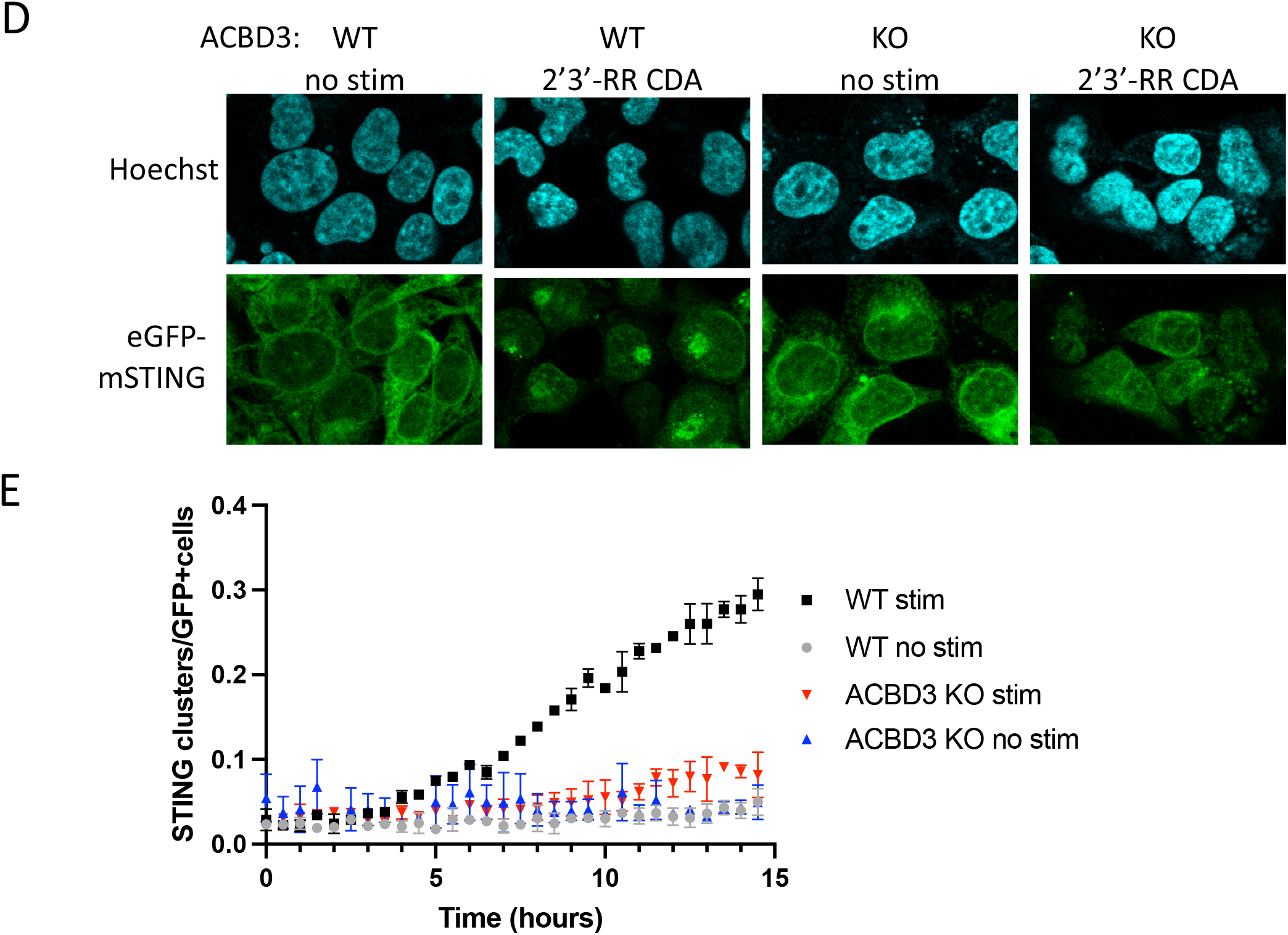
ACBD3 is important for STING phosphorylation, clustering and relocalization. a. Normalized [^32^P]2′3′-cGAMP (cGAMP) and [^32^P]3′3′-c-di-AMP (CDA) uptake after 1 h by THP-1 monocytes transduced with a non-targeting control CRISPRi gRNA, or SLC19A1 or ACBD3 CRISPRi gRNA. Mean ± SEM of *n* = 3 biological replicates are shown. b. Immunoblot analysis of protein expression and phosphorylation in THP-1 cells expressing indicated CRISPRi gRNAs. Cells were stimulated for 2h with 2’3’-RR CDA (5μg/ml) or left unstimulated. p-STING: STING phosphorylated on S366. Representative images of *n* = 2 biological replicates are shown. c. Immunofluorescence of 293T cells expressing STING-HA and a control CRISPRi gRNA or an ACBD3-targeting CRISPRi gRNA. Cells were stimulated for 2h with 2’3’-RR CDA (STING agonist) and stained for HA or ACBD3. Representative images of *n* = 2 biological replicates are shown. d. Immunofluorescence live-cell imaging eGFP-tagged mouse-STING in control 293T cells or 293T cells lacking ACBD3 (KO) stimulated with 2’3-RR CDA for 11.5h. Representative images of *n* = 2 biological replicates are shown. e. The number of STING clusters/eGFP+ cells shown in d was quantified over time using the ‘particle analysis’ function of ImageJ. Mean ± SEM of *n* = 2 biological replicates are shown.

STING phosphorylation requires STING trafficking from the ER to the trans-Golgi network (TGN), where STING palmitoylation promotes the formation of activation clusters needed for downstream signalling^23^. Indeed, STING is recruited to perinuclear clusters in HA-STING expressing 293T cells and THP-1 cells upon STING activation (Figure 2C and S3). These clusters colocalized with the Golgi-resident protein ACBD3. Upon stimulation with 2’3’-RR-CDA in ACBD3-depleted cells, STING did not cluster, and its localization appeared similar to that of unstimulated cells, presumably the ER. Similarly, live-cell imaging of eGFP-tagged mouse STING (eGFP-mSTING) showed cluster formation in control 293T cells, but not in cells lacking ACBD3 (Figure 2D and 2E). Thus, ACBD3 is required for the relocalization of STING upon activation.

ACBD3 binds and recruits PI4KB to the TGN^24^. In line with this, ACBD3 colocalized to TGN46 (a Golgi marker) and PI4KB-enriched perinuclear clusters in unstimulated THP-1 cells (Figure 3A and 3B). In ACBD3 depleted cells, perinuclear PI4KB clusters were absent (Figure 3B). Similarly, PI4KB clustered with PI4P in control cells, whereas perinuclear clustering of PI4P and PI4KB was lost in ACBD3 depleted cells (Figure 3C). PI4P and p-STING colocalized in perinuclear clusters upon STING activation of control cells, but not upon ACBD3 depletion (Figure 3D).

**Figure 3.**
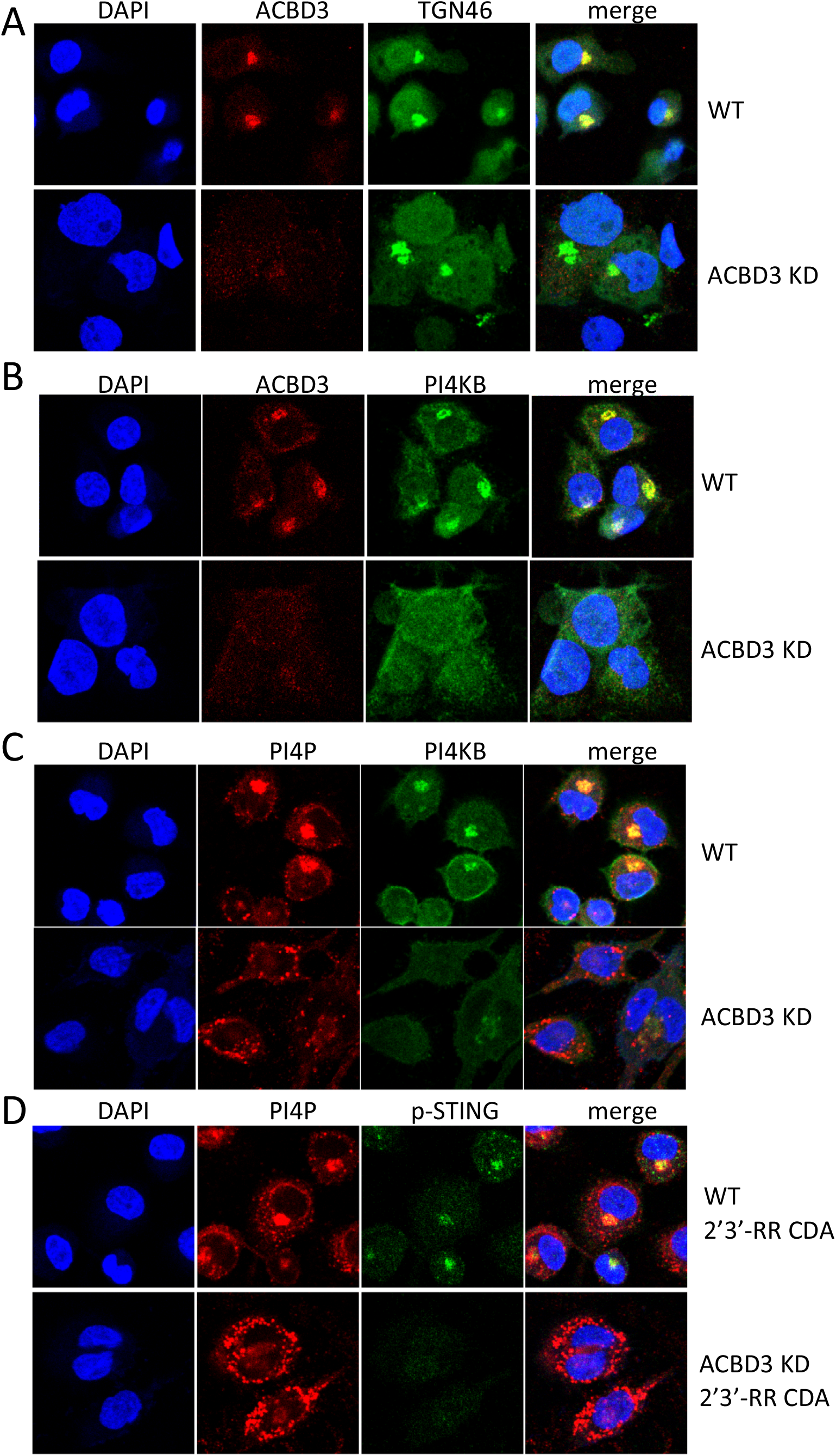
STING localizes to PI4P-rich environments a-d. Immunofluorescence images of THP-1 cells expressing control or ACBD3 CRISPRi gRNAs stained for the indicated proteins. During cell culturing, THP-1 cells were treated with PMA to make them adherent to the coverslips. In panel d, THP-1 cells were stimulated for 2h with 5ug/ml 2’3’-RR CDA prior to staining. a-d. Representative images of at least *n* = 2 biological replicates are shown.

The role of PI4KB in STING activation was supported by our genome wide screens for STING-associated factors, where we found that PI4KB was a significant hit in the screen for genes whose depletion resulted in weaker activation of the pathway^20^. We confirmed the role of PI4KB in STING pathway activation by treating cells with the PI4KB inhibitor BF738735^25^, which reduced STING pathway activation (Figure 4A). Furthermore, when we depleted PI4KB expression using CRISPRi gRNAs we observed that gRNAs that depleted progressively more *Pi4kb* mRNA led to progressively greater deficiency in reporter activation after stimulation with STING agonist (Figure 4B).

**Figure 4.**
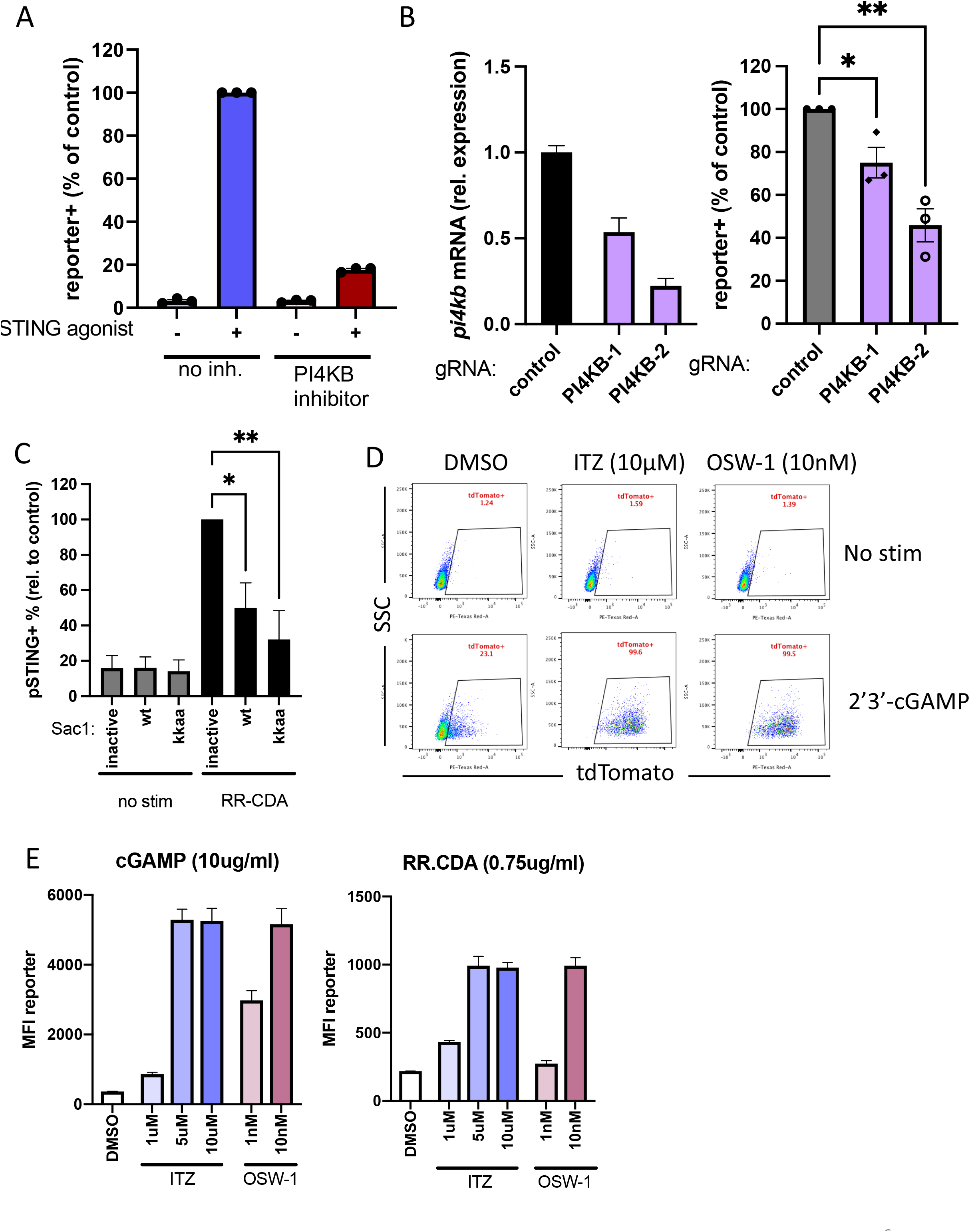
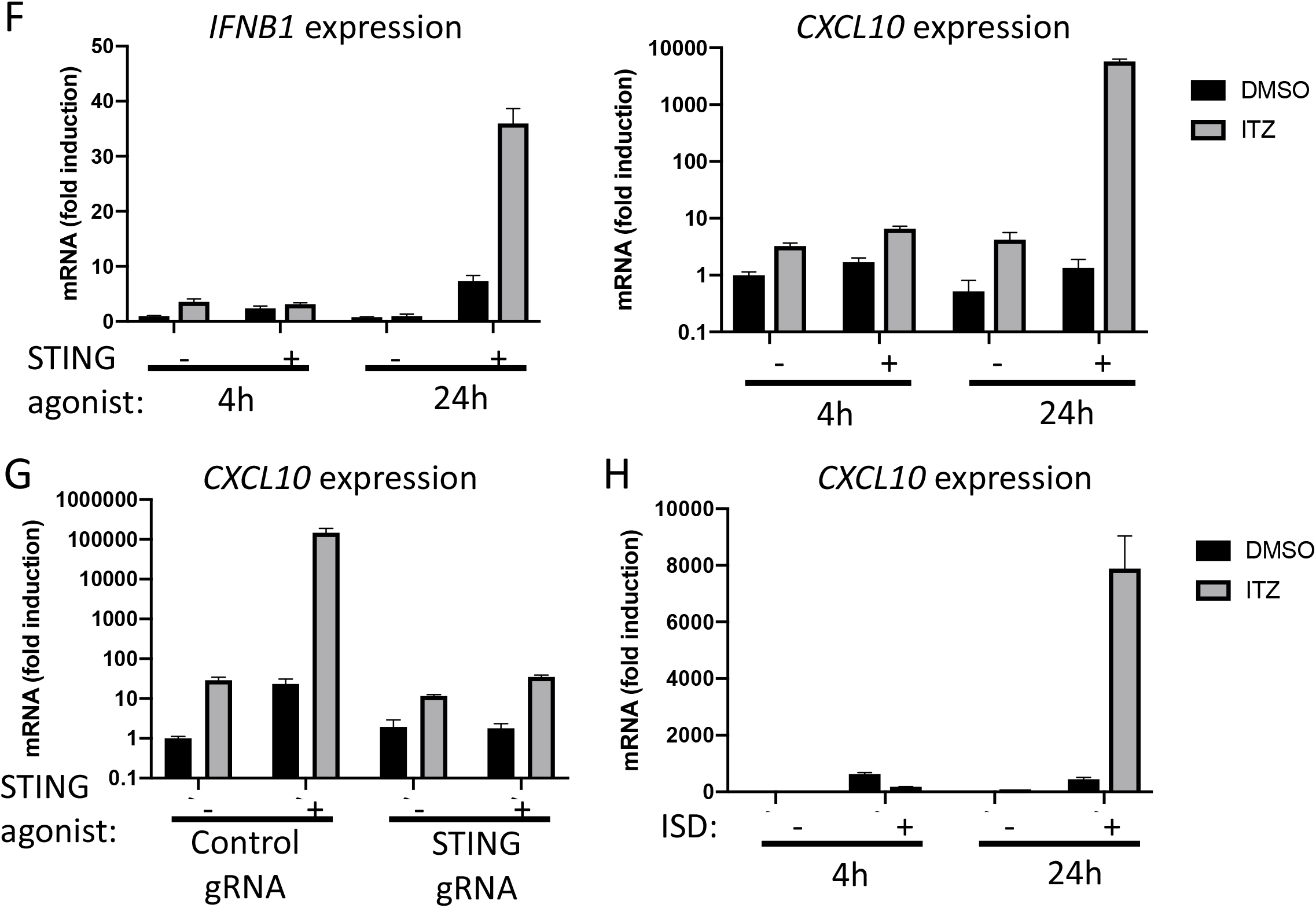
PI4KB and OSBP inhibition have opposite effects on STING activation a. THP-1 cells were pre-incubated with the PI4KB inhibitor BF738735 (10uM) and subsequently stimulated with the STING agonist 2’3’-RR CDA (1.67ug/ml). After 20-24h, tdTomato-reporter expression was quantified by flow cytometry. b. (left panel) *PI4KB* mRNA expression levels in THP-1 cells expressing a control gRNA or gRNAs targeting PI4KB. Mean ± SEM of *n* = 3 technical replicates are shown. (right panel) Cells were stimulated with 2’3’-RR CDA for 20-24h and tdTomato reporter expression was quantified by flow cytometry. Mean ± SEM of *n* = 3 biological replicates are shown. c. 293T cells expressing eGFP-mSTING were transfected with a phosphatase-dead Sac1 (inactive), active Sac1 wt, or Sac1-kkaa mutant and stimulated or not with 2’3’-RR CDA (RR-CDA) for 8h. After stimulation, cells were stained for phospho-STING and analyzed by flow cytometry. Mean ± SEM of *n* = 3 biological replicates are shown. d. THP-1 cells were preincubated with DMSO or the OSBP inhibitors itraconazole (ITZ) or OSW-1 and stimulated with 2’3’-cGAMP (7μg/ml) or left untreated. Reporter expression was quantified by flow cytometry 20h after stimulation. Representative images of *n* = 3 biological replicates are shown. e. Mean fluorescence intensity of tdTomato reporter in THP-1 cells stimulated with STING agonists for 20h-24h in the presence of DMSO, itraconazole (ITZ), or OSW-1. f. *IFNB1* or *CXCL10* mRNA levels in THP-1 cells pre-treated with DMSO or itraconazole (ITZ) for 1h and stimulated with 2’3’-RR CDA (RR-CDA) for 4 or 24h. g. CXCL10 mRNA levels in THP-1 cells expressing a control gRNA or STING gRNA (STING knockdown; KD) treated as in e and measured after 24h. h. CXCL10 mRNA levels in THP-1 pre-treated as in e and transfected with VACV-70 dsDNA for 4h or 24 e-h. Mean ± SEM of *n* = 3 biological replicates are shown.

Upon shuttling to the ER, PI4P is ultimately hydrolysed by the phosphatase Sac1. Overexpression of wild type Sac\1 (Sac1-wt) or a Sac1- K583A K585A double mutant that localizes to the TGN (Sac1-kkaa) effectively depletes PI4P at the TGN, in contrast to a phosphatase-dead version of Sac1^26^. In 293T cells expressing eGFP-mSTING, overexpression of Sac1-wt or the Sac1-kkaa mutant significantly impaired STING activation by CDNs compared to overexpression of the phosphatase-dead mutant of Sac1 (Figure 4C). Taken together, these results suggest that PI4P levels at the TGN are important for proper STING activation.

Another protein that regulates PI4P levels at the TGN is oxysterol binding protein (OSBP). This protein shuttles PI4P from the TGN to the ER and/or to lysosomes in exchange for cholesterol. Inhibition of OSBP by the plant-extract OSW-1 or the FDA-approved drug itraconazole results in the accumulation of PI4P at the TGN^27,28^. Considering that reducing PI4P at the TGN by targeting ACBD3 or PI4KB hampered STING activation, we hypothesized that increasing PI4P concentrations at the TGN by targeting or inhibiting OSBP might result in enhanced STING signalling. We tested this by combining itraconazole or OSW-1, which did not by themselves activate the STING pathway, with a limiting dose of RR-CDA or 2’3’-cGAMP that by themselves induced only a small response in THP-1 cells. Combining the OSBP inhibitors with limiting doses of STING agonist resulted in a dramatic increase in reporter activation (Figure 4D and 4E). Synergistic pathway activation was also observed when we tested induction of the endogenous transcripts *CXCL10* and *IFNB1* (Figure 4F). The effect was dependent on STING expression (Figure 4G). Finally, immune activation resulting from transfection of DNA, which induces endogenous production of 2’3’-cGAMP, was also enhanced in the presence of itraconazole (Figure 4H). Overall, these results indicate that drugs that modulate PI4P levels may have promise for either boosting or restraining STING pathway activation.

To further dissect the mechanism by which OSBP inhibition increases STING activation, we tested the phosphorylation status of STING at different time points after stimulation (Figure 5A). Relative to the results after stimulation with 2’3’ cGAMP alone, the addition of itraconazole resulted in a clear increase in STING phosphorylation at 4h and 8h, but had little or no effect at 2h. OSBP inhibition in the absence of STING agonists did not promote STING phosphorylation (Figure S4A). When combined with 2’3’ cGAMP, OSBP inhibitors also enhanced the phosphorylation of TBK-1 and IRF3, and the degradation of IκBα, the latter a marker of NF-κB activation (Figure S4B). Furthermore, stimulating cells with limiting amounts of 2’3’-cGAMP in the presence of OSBP inhibitors promoted eGFP-mSTING clustering (Figure 5B and 5C). The pronounced STING activation induced by itraconazole did not result from an increase in 2’3’-cGAMP taken up from the extracellular environment (Figure S4C).

**Figure 5.**
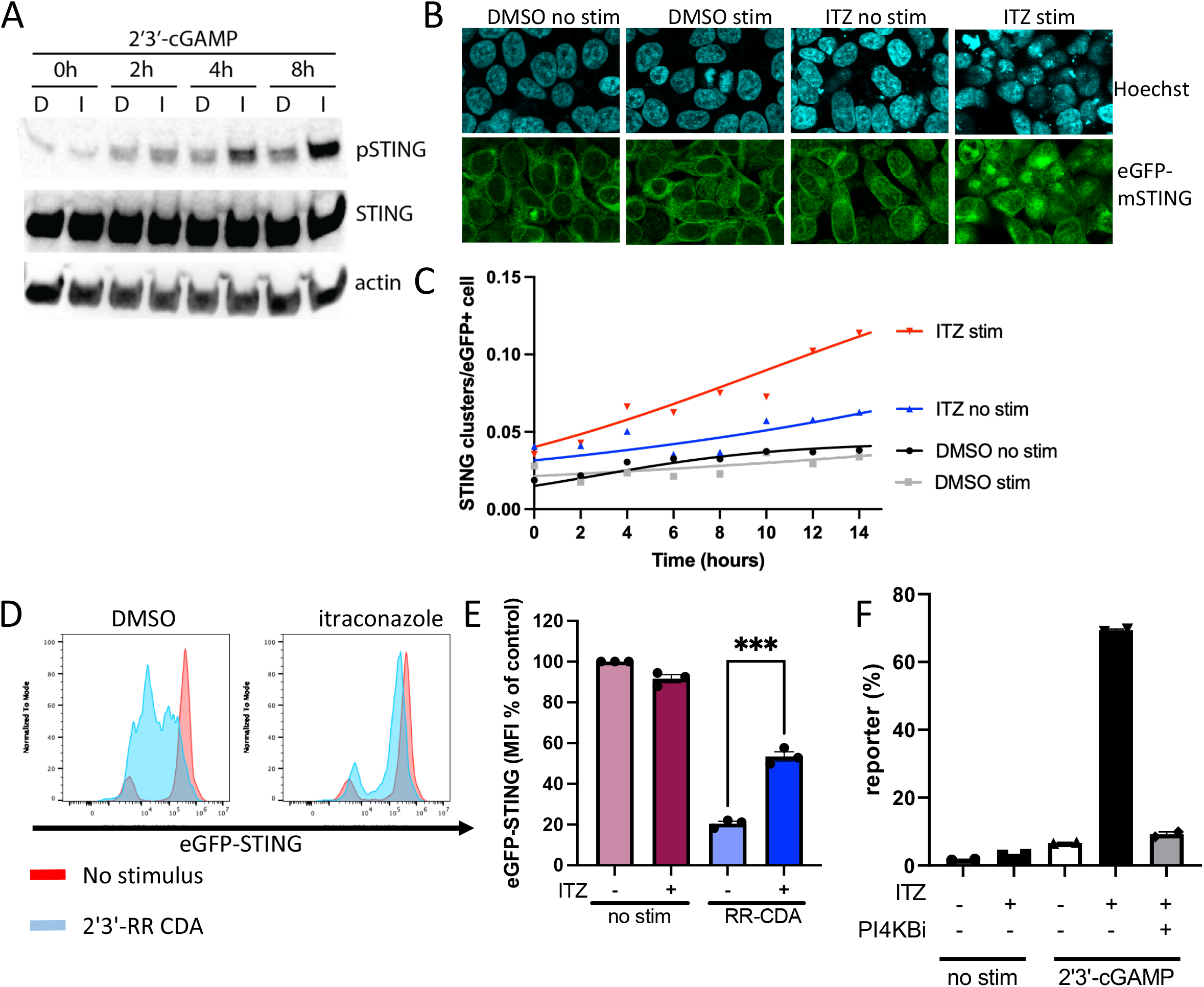
OSBP inhibition increases STING activation and decreases STING degradation a. Immunoblot analysis of the indicated (phosphorylated) proteins expressed by THP-1 cells. Cells were pre-treated with DMSO (D) or itraconazole (I) for 1h and stimulated with 2’3’-cGAMP (20ug/ml) for the indicated time points. Representative images of *n* = 3 biological experiments are shown. b. Immunofluorescence live-cell imaging eGFP-tagged mouse-STING in 293T cells pre-treated for 1h with DMSO or itraconazole (ITZ) and stimulated with 2’3’-cGAMP for 15h. Representative images of *n* = 2 biological experiments are shown. c. The number of STING clusters/GFP+ cells shown in c was quantified over time using the ‘particle analysis’ function of ImageJ. Representative quantification of *n* = 2 biological experiments is shown. d. Expression of eGFP-tagged mouse STING in THP-1 cells pre-treated with DMSO or itraconazole for 1h and stimulated with 2’3’-RR CDA (10μg/ml) or left unstimulated for 20h. Representative image of *n* = 3 biological replicates is shown. e. Quantification of eGFP-STING in THP-1 cells shown in e. Mean ± SEM of *n* = 3 biological replicates are shown. f. tdTomato reporter expression of THP-1 cells pre-treated with the PI4KB inhibitor (PI4KBi) BF738735 for 1h followed by pre-treatment with itraconazole (ITZ) for 1h and stimulation with 2’3’-cGAMP for 20h. Mean ± SEM of *n* = 2 biological replicates are shown.

After activation, STING traffics to the endolysosomal compartment for degradation^29^. To quantify STING degradation, eGFP-mSTING expression was measured 20h after activation in the presence of DMSO or itraconazole (Figure 5D). Upon activation, eGFP-mSTING was degraded in control cells, but degradation was significantly reduced in the presence of itraconazole (Figure 5E).

Finally, we investigated the role of PI4KB in STING activation by OSBP inhibitors using the PI4KB inhibitor BF738735. PI4KB inhibition completely reverted the amplifying effect of itraconazole on STING activation (Figure 5F). Thus, PI4P production at the TGN is critical for STING activation and can be targeted to promote or dampen the STING-induced immune response.

## Discussion

Here, we show for the first time that the phospholipid PI4P plays a critical role in the STING-induced immune response. Upon activation, STING trafficked to PI4P-positive structures that were regulated by PI4KB and ACBD3, the latter being one of the top hits in our screen for genes required for STING activation^30^. Depleting ACBD3 blocked STING activation by various cyclic dinucleotides (CDNs) and other STING agonists by preventing STING trafficking to the TGN. STING trafficking from the ER to the TGN is critical for downstream immune activation. At the TGN, STING forms oligomers that interact with TBK1^31^, leading to phosphorylation of TBK1, STING and IRF3^13^. After immune activation, STING is degraded by the endolysosomal system or transported back to the ER to terminate immune signalling^32^. The mechanisms regulating trafficking and retention of STING in the Golgi, and subsequent switch to Golgi exit are not known^13^. We show that the intensity of STING activation and its subsequent degradation depends on proteins that affect PI4P levels at the TGN, including PI4KB, Sac1, OSBP, and ACBD3.

ACBD3 is a multifunctional protein involved in various cellular processes, including recruitment of PI4KB to the TGN^33^, hormone-induced steroid formation at mitochondria by binding to PKA^34^, and iron uptake by the divalent metal transporter DMT1^35^. As PI4KB was also a hit in our primary screen for STING activation, but not any of the other known ACBD3 binding partners, we focussed on the role of ACBD3 and PI4KB in STING activation. As reported previously^36^, we observed that ACBD3 depletion dramatically altered the intracellular distribution of PI4KB and PI4P. The role of PI4P in STING activation was established by reducing PI4P production at the TGN either by inhibiting PI4KB or by increasing PI4P hydrolysis by Sac1. Conversely, increasing PI4P levels by inhibiting the lipid transfer protein OSBP dramatically enhanced STING activation. Thus, PI4P levels at the TGN dictate the intensity of the STING-induced immune response.

PI4P at the TGN is produced by PI4KB and PI4K2A, although these kinases act in different subregions^27^. Targeting PI4KB strongly reduced STING activation, suggesting that PI4KB produces the main pool of PI4P needed for STING activation. Residual STING activity observed upon inhibition or depletion of PI4KB is likely due to the activity of other kinases, including PI4K2A.

How PI4P affects STING activation remains unknown. PI4P lipids can anchor various proteins to the Golgi via their PI4P-interacting domains, such as pleckstrin homology (PH) domains. Binding is partly driven by electrostatic interactions between the inositol head-group of PI4P and cationic residues in PH domains^37^. Recently, the purified C-terminal domain of STING was shown to bind PI4P lipids^38^. Although STING shows no homology to other known PI4P-binding domains, computational modelling of STING in an active conformation pointed to a patch of basic amino acids in close proximity to the transmembrane helices of STING that can accommodate PI4P^38^. Although more data is needed to confirm STING binding to PI4P in cells, such an interaction may regulate the duration of signalling by anchoring ligand-bound STING to the TGN. Increasing PI4P levels at the TGN (e.g. by blocking OSBP) may thus increase STING-mediated signalling by prolonging its retention at the TGN, thereby delaying subsequent degradation. Similarly, degradation-resistant STING mutants show improved downstream signalling upon activation.^29^ Besides directly interacting with STING, PI4P may promote STING trafficking by facilitating the general process of ER-to-Golgi transport of proteins. In yeast, for example, COP-II vesicle fusion depends on cis-Golgi-localized PI4P^39^. In mammalian cells, however, the role of PI4P in COP-II vesicle transport remains to be elucidated.

OSBP shuttles TGN-localized PI4P to the ER in exchange for cholesterol, which moves in the opposite direction. Thus, OSBP inhibition not only increases PI4P at the TGN, but at the same time increases cholesterol levels at the ER membrane^40^. The subcellular distribution of cholesterol may also affect STING activation. For example, STING is constitutively active in cells lacking the lysosomal cholesterol transporter NPC1 due to a reduction in ER-cholesterol^41^. In line with this, STING was shown to be activated upon a decrease in ER-localized cholesterol in another study^42^. Although a build-up of cholesterol in the ER upon OSBP inhibition may dampen STING activation, we observed a strong increase in STING activation, indicating that the accumulation of PI4P is a dominant factor in regulating STING activation. This is in line with our observations that inhibiting PI4P production nullifies the boosting effect of OSBP inhibitors. Our results also indicate that the previously documented increase in STING activation upon ER-cholesterol depletion may be caused by PI4P accumulation at the TGN, as ER-cholesterol depletion prevents PI4P shuttling by OSBP, and has thus a similar effect as OSBP inhibition^27^.

Itraconazole is an established antifungal and is being evaluated as an anticancer drug^43^. Itraconazole has, apart from OSBP^28^, several targets including the Hedgehog pathway^44^, VEGF2^45^, VDAC1^46^ and NPC1^47^. Knockout of the lysosomal protein NPC1 results in tonic STING activation due to the depletion of ER cholesterol, which causes relocation of the cholesterol sensor SREBP2 and STING to the Golgi, and by preventing lysosomal degradation of STING, thereby boosting immune activation^41^. NPC1 knockout furthermore results in PI4P accumulation at the TGN, and may thus further promote STING activation^48^. Although we did not observe tonic STING activation in itraconazole-treated cells, inhibition of NPC1 may partially explain the boosting effect of itraconazole. In contrast, the structurally unrelated OSBP-inhibitor OSW-1 is highly specific for OSBP, and interacts with OSBP in the nanomolar range via a binding site that is different from itraconazole^49^. OSW-1 is not known to inhibit NPC1 or any other target of itraconazole. Thus, the effect of OSW-1 or itraconazole on STING activation is likely via OSBP inhibition. Supporting this, NPC1, SREBP2, or other cholesterol-regulating factors were not identified as hits in our genome-wide screens for STING regulators^20^.

Enhancing STING activation by itraconazole or other OSBP inhibitors has therapeutic potential by promoting the immune response to virus-infected or cancer cells. In cancer cells, accumulation of cytosolic DNA can activate the cGAS/STING pathway and promote tumour clearance, although some cancer cells epigenetically silence STING or express STING mutants with reduced activity^50^. In these cases, increasing Golgi PI4P levels (e.g. via OSBP inhibition) may improve the endogenous immune response. Furthermore, treatment of tumours with DNA damaging agents^51^ or irradiation^52,53^ can invoke STING activation, and this may be enhanced upon OSBP inhibition. OSBP inhibitors may also improve the antitumor effects of STING agonists used therapeutically. In line with this proposal, intratumoural injection of cGAMP in combination with bafilomycin A1, which prevented lysosomal degradation of STING, dramatically improved tumour cell clearance *in vivo*^29^. STING is also frequently targeted by viruses in infected cells, thereby dampening the innate immune response^54^. In that instance, inadequate STING activation may also be restored by treatment with itraconazole or other OSBP inhibitors.

In conclusion, we found that STING activation is controlled by PI4P. Targeting this pathway by (FDA-approved) compounds, opens up new avenues for therapies that depend on proper STING activation.

## Acknowledgements

We thank Richard Wubbolts and Esther van t Veld (Centre for Cell Imaging, Faculty of Veterinary Medicine, Utrecht University) and Denise Schichnes (the Biological imaging facility, UC Berkeley) for support with the fluorescent microscopy experiments, Ger Arkesteijn (Flow Cytometry Facility, Faculty of Veterinary Medicine, Utrecht University), Hector Nolla and Alma Valeros (Flow Cytometry Facility, UC Berkeley) for technical support with flow cytometry experiments, Matthew Shair and Peter Mayinger for sharing reagents, Lily Zhang (UC Berkeley) for lab support, and the members of the Raulet and van Kuppeveld labs for helpful discussions. R.D.L. was supported by the Cancer Research Institute Irvington Postdoctoral Fellowship and a Marie Skłodowska-Curie Individual Fellowship. The research was supported by grant AI113041 from the US National Institutes of Health to DHR. J.J.W. was supported by NIH grant R21-AI137758. S.A.Z. was supported by the University of Washington/Fred Hutchinson Cancer Research Center Viral Pathogenesis Training Program (AI083203), the University of Washington Medical Scientist Training Program (GM007266) and the Seattle ARCS foundation.

## Conflicts of Interest

DHR is a cofounder and SAB member of Dragonfly Therapeutics, and a member of the SAB of Vivere Pharmaceuticals.

## Materials and Methods

### Cell lines

All cell lines were cultured at 37 °C in humidified atmosphere containing 5% CO_2_ with medium supplemented with 100 U ml^−1^ penicillin, 100 μg ml^−1^ streptomycin, 0.2 mg ml^−1^ glutamine, 10 μg ml^−1^ gentamycin sulfate, 20 mM HEPES and 10% heat-inactivated FCS. THP-1 cells were cultured in RPMI medium, and 293T, 293T transfected with human STING (293T+hSTING) were cultured in DMEM medium. THP-1 and 293T cells were from existing stocks in the laboratory. The 293T+hSTING cells were generated as described previously^14^.

### Antibodies and reagents

The following antibodies were obtained from Cell Signaling Technology: rabbit-anti-human TBK1 monoclonal (clone D1B4; 1:500 for immunoblot), rabbit-anti-human p-TBK1 monoclonal (clone D52C2; 1:1,000 for immunoblot), rabbit-anti-human STING monoclonal (clone D2P2F; 1:2,000 for immunoblot), rabbit-anti-human p-STING monoclonal (clone D7C3S; 1:1,000 for immunoblot and 1:800 for flow cytometry), rabbit-anti-human p-IRF3 monoclonal (clone 4D4G, 1:1,000 for immunoblot) rabbit-anti-IκBalpha (clone 9242S, used 1:500 for immunoblot). Antibodies obtained from LI-COR Biosciences: goat-anti-mouse IgG IRDye 680RD conjugated (cat. no. 926-68070; used at 1:5,000), donkey-anti-rabbit IgG IRDye 800CW conjugated (cat. no. 926-32213; used at 1:5,000), donkey-anti-rabbit IgG IRDye 680RD (cat. no. 926-68073; used at 1:5,000). Other antibodies: rabbit-anti-human IRF3 monoclonal (Abcam, cat. no. EP2419Y; 1:2,000 for immunoblot), mouse-anti-human transferrin receptor monoclonal (Thermo Fischer Scientific, clone H68.4; 1:1,000 for immunoblot), mouse-anti-Actin (Sigma cat. no A5441, 1:5,000 for immunoblot), rabbit-anti PI4KB (FineTest cat.no. FNab06427, 1:100 for immunofluorescence), mouse IgM-anti PI4P (Echelon Biosciences Z-P004; 1:100 for immunofluorescence), rabbit-anti TGN46 (Novus Biologicals cat. no. NBP1-49643, 1:400 for immunofluorescence), mouse-anti ACBD3 (Sigma cat. no. Sigma WH0064746M1, 1:100 for immunofluorescence and flow cytometry, 1:1000 for immunoblot), rat-anti-HA (clone 3F10, Roche cat. no. 11867423001, 1:500 for immunofluorescence). Secondary antibodies from Invitrogen: Goat-anti-mouse AlexaFluor 488-conjugated (cat. no. A11001), goat-anti-rat Alexa Fluor 568-conjugated (cat. no A11011), goat-anti-mouse-IgM Alexa Fluro 568-conjugated (cat. no. A21043), donkey-anti-rabbit AlexaFluor 647-conjugated (cat. no. A31573), donkey-anti-mouse AlexaFluor 568-conjugated (cat. no. A10037), and donkey-anti-mouse AlexaFluor 647-conjugated (cat. no. A10037).

Reagents used include itraconazole (Santa Cruz Biotechnology cat. no. sc-205724A), OSW-1 (a kind gift from M. Shair, Harvard University), BF738735 (Tocris cat.no. 6246/10), polybrene (EMD Millipore, cat. no. TR1003G), diABZI (Invivogen cat. no. tlrl-diabzi), 3′3′-cyclic-di-AMP (3′3′ CDA) (Invivogen, cat. no. tlrl-nacda), 2′3′-RR c-di-AMP (2′3′-RR-S2 CDA) (Invivogen cat. no. tlrl-nacda2r), 2′3′- cyclic-di-GMP-AMP (2′3′-cGAMP) (Invivogen cat. no. tlrl-nacga23), and human IFN-β (PeproTech, cat. no. 300-02B). Antibiotic selection was carried out with puromycin (Sigma-Aldrich, cat. no. P8833, at 2 μg ml^−1^), blasticidin (Invivogen, cat. no. ant-bl-1, at 10 μg ml^−1^), and zeocin (Invivogen, cat. no. ant-zn-1, at 200 μg ml^−1^).

### Plasmids and expression

The lentiviral vector encoding the tdTomato reporter gene driven by the ISREs and the minimal mouse IFN-β promoter was generated as described previously^30^. For rescue and overexpression, *ACBD3* was cloned into a dual promoter lentiviral vector co-expressing the blasticidin resistance gene and the fluorescent gene mAmetrine^55^. For over-expression of eGFP-coupled to the N terminus of mouse STING (mSTING) via a linker sequence (amino acid sequence GAGAKLGTELGS), the fusion construct was cloned using Gibson assembly into a dual promoter lentiviral vector co-expressing the blasticidin gene. For CRISPR interference (CRISPRi)-mediated depletions, cells were transduced with a lentiviral dCas9-HA-BFP-KRAB-NLS expression vector (Addgene plasmid no.102244).

For gene depletions using individual CRISPRi gRNAs, top enriched gRNAs (Supplementary Table 1) from the screen for STING activation were cloned into the same expression plasmid used for the gRNA library (pCRISPRia-v2, Addgene plasmid no. 84832, a gift from J. Weissman). The lentiviral gRNA plasmid co-expressed a puromycin resistance gene and blue fluorescence protein (BFP) via a T2A ribosomal skipping sequence controlled by the human EF1A promoter. Conventional CRISPR gRNAs (see Supplementary Table 1) were cloned into a puromycin-selectable lentiviral CRISPR–Cas9 vector, as described previously^56^. *Sac1* wild type (a generous gift from Peter Mayinger (OHSU), the catalytically-inactive C389S mutant, or the Golgi-directed K583A K585A double mutant (kkaa) was fused to the N-terminus of the fluorescent gene mTurquoise2 via a linker sequence (encoding MTSKSGGGGSGGGG) and cloned using NEBuilder Hifi DNA assembly (New England Biolabs) into a dual promoter lentiviral vector co-expressing a puromycin resistance gene. Lentivirus was produced by transfecting lentiviral plasmids and second generation packaging and polymerase plasmids into 293T cells, as described previously^30^. *Sac1*-encoding lentiviral plasmids were not used for lentivirus production, but directly transfected into cells for phospho-STING analysis using lipofectamine 2000 (Invitrogen), as lentiviral over-expression of Sac1 was toxic to cells.

### CDN and IFN-β stimulation reporter assays

Stimulation with CDNs or IFN-β were performed as described previously^30^. Briefly, the day before stimulation, cells were seeded to 0.5 × 10^6^ cells per ml. Cells were stimulated with CDNs or IFN-β in 96-well plates using 30,000 cells per well in 150 μl medium. After 18–24 h, cells were transferred to a 96-well plate and tdTomato expression was measured by flow cytometry using a high-throughput plate reader on a BD LSR Fortessa or a Beckman Coulter Cytoflex. For stimulations in the presence of itraconazole, OSW-1 and/or BF738735, cells were incubated with compounds or DMSO as vehicle 1h before stimulations with CDNs or IFN-β. 18–24h after stimulation, tdTomato reporter expression was quantified by flow cytometry using a high-throughput plate reader on a BD LSR Fortessa or a Beckman Coulter Cytoflex.

### Production of *ACBD3*-knockout cell lines

As an alternative approach to corroborate the role of *ACBD3* in CDN responses, *ACBD3* was targeted in THP-1 or 293T cells using the conventional CRISPR–Cas9 system. THP-1 cells were transduced and 293T cells were transfected with a CRISPR–Cas9 lentiviral plasmid encoding a control gRNA or a gRNA targeting *ACBD3* (see Supplementary Table 1). After transduction/transfection cells were selected using puromycin for two days and single-cell cloned by limited dilution (100 cells diluted in 50ml, and plated on 96W plates using 200 μl/well). Control cells and *ACBD3*-targeted cells were selected that had comparable forward and side scatter by flow cytometry analysis, and *ACBD3*-knockout cells were screened by measuring intracellular ACBD3-expression by flow cytometry.

### CDN uptake

The production and uptake of [^32^P]2’3’-cGAMP and [^32^P]3’3’-CDA was performed as described previously^30^. For the uptake of 2’3’-cGAMP in cells treated with itraconazole (Figure S4C), cells were pre-treated for 1h with itraconazole, and subsequently stimulated for 8h in the presence of 2’3’-cGAMP (20 μg/ml). After stimulation, cells were washed twice with ice cold PBS and pellets were lysed in H_2_O. 2’3’-cGAMP levels in cell lysates were tested using a 2’3’-cGAMP ELISA kit (Cayman Chemical cat. no. 501700) according to the manufacturer’s recommendations.

### Stimulation for RT-qPCR or immunoblotting

The day before stimulation, cells were seeded to 0.5 × 10^6^ cells per ml. Cells were stimulated with CDNs or transfected with VACV-70 immunostimulatory DNA using lipofectamine 2000 using 0.5 × 10^6^ cells per well in 500 μl medium. For stimulations in the presence of itraconazole or OSW-1, cells were incubated with compounds or DMSO as vehicle 1h before stimulations After stimulations, cells were further processed (see RT-qPCR and immunoblotting).

### RT–qPCR

Cells were collected and washed in ice-cold PBS. Cells were transferred to RNase-free microcentrifuge tubes and RNA was isolated using the RNeasy mini kit (Qiagen, cat. no. 74104) including a DNase I step (Qiagen, cat. no. 79254). RNA concentration was measured by NanoDrop (Thermo Fischer), and 1 μg of RNA was used as input for cDNA synthesis using the iScript cDNA synthesis kit (Bio-rad, cat. no. 1708890) or Superscript III (Invitrogen cat. no. 18080) using random hexamers. cDNA was diluted to 20 ng μl^−1^ and 2.5 μl per reaction was used as input for the qPCR reaction. qPCR reactions were set up using SSOFast EvaGreen Supermix (Bio-Rad, cat. no. 1725200) or Fast SYBR Green master mix (Applied Biosystems cat. no. 4385612) according to the manufacturer’s recommendations, using 500 nM of each primer and following cycling conditions on a Bio-Rad C1000 Thermal Cycler or Roche Lightcycler 480 II: 2 min at 98 °C, 40 repeats of 2 s at 98°C and 5 s at 55°C. Primers used to amplify the PCR-products specific for the human genes *HPRT1, YWHAZ, IFNB1, IL-6, CXCL10*, and *PI4KB* are listed in Supplementary Table 2. The housekeeping genes *HPRT1* and *YWHAZ* served as endogenous controls for cDNA samples.

### Cell lysis and immunoblotting

For protein detection by immunoblotting, cells were washed with PBS and lysed in RIPA buffer (25 mM Tris-HCl pH 7.5, 150 mM NaCl, 1 mM EDTA, 1% NP-40 and 0.1% SDS) including cOmplete ULTRA protease inhibitors (Sigma-Aldrich cat. no. 05892791001), phosphatase inhibitors (Biomake, cat. no. B15001) and 50 mM DTT. Cell lysates were mixed with 4× NuPage LDS sample buffer (Invitrogen cat. no. NP0007), pulse sonicated and incubated at 75 °C for 5 min. Lysates were loaded onto Bolt 4–12% Bis-Tris Plus SDS–PAGE gels (Invitrogen cat. no. NW04125BOX). Proteins separated by SDS–PAGE were transferred onto Immobilon-FL PVDF membranes (EMD Millipore) at 100 V for 1 h at 4 °C. Membranes were blocked in 4% NFM, and probed in 1% NFM overnight at 4 °C with primary antibody. Membranes were subsequently washed three times in 1× TBS including Tween 20 (0.05%) (TBS-T) and probed with secondary antibody for 1 h at room temperature while protected from light. Membranes were washed two times in TBS-T, once in TBS and blots were imaged using an Odyssey CLx System (LI-COR).

### Intracellular phospho-STING stainings upon *Sac1* transfection

293T cells stably expressing eGFP-mSTING were transfected with plasmids encoding Sac1-wt and mutants fused to the fluorescent reporter mTurquoise2. Twenty-four hours after transfection, cells were stimulated for 8h with 2’3’-RR CDA. After stimulation, cells were washed and blocked using Trustain Fc receptor blocking solution (Biolegend cat. no. 422302) for 10min at RT. Cells were fixed in 2% formaldehyde in PBS for 10min at 4 °C. Cells were permeabilized in perm/wash buffer (BD Biosciences cat. no. 554714) for 15min at 4 °C. Cells were incubated with primary antibody in perm/wash buffer for 30min at 4 °C, washed, and incubated in secondary antibody for 30min at 4 °C. Cells were washed and analyzed by flow cytometry (Beckman Coulter Cytoflex).

### Confocal microscopy

The day before seeding onto microscopy slides, cells were seeded to 0.5 × 10^6^ cells per ml. For live-cell imaging, 293T cells were reseeded onto a Ibidi 4-well chambers (Ibidi cat. no. 80416) treated the day before with 5μg/ml fibronectin (Sigma Aldrich cat. no. F1141). Cells were allowed to recover for 2 days and used for live cell imaging in a humidified, temperature and CO2-controlled chamber using a Nikon A1R confocal microscope. For fixed-sample confocal microscopy, 293T or THP-1 cells were reseeded onto an Ibidi μ slide 18 well (cat.no. 81826) treated the day before with 5μg/ml fibronectin. 293T cells were allowed to recover for 2 days prior to stimulation and staining. Prior to stimulation and staining of THP-1 cells, cells were treated overnight with 30 ng/ml PMA (Sigma Aldrich cat. no. P1585) followed by overnight recovery in PMA-free medium. Cells were stimulated with CDNs for the indicated time points and fixed using 2% formaldehyde for 15min at RT. Samples were incubated with 50mM NH_4_Cl in PBS for 10min at RT and permeabilized with 0.2% Triton X100 in PBS for 15min at RT. Samples were blocked in 3% BSA and 0.2% Triton X-100 in PBS for 45min at RT. Cells were washed incubated with indicated primary antibodies for 1h at RT, washed, and incubated in secondary antibodies for 1h at RT in 0.3% BSA and 0.02% Triton X-100. Cells were washed and kept in PBS + DAPI at 4 °C until imaging on a Nikon A1R confocal microcope. For PI4P staining, after fixing, cells were permeabilized with 20 μM digitonin in buffer A (20mM Pipes, pH 6.8, 137mM NaCl, 2.7mM KCl). Cells were blocked using 5% normal goat serum (NGS) and 50mM NH_4_Cl in buffer A for 45min at RT. Cells were incubated in primary antibodies in buffer A supplemented with 5% NGS for 1h at RT, washed, and incubated in secondary antibodies in buffer A supplemented with 5% NGS for 1h at RT. Cells were incubated in 2% formaldehyde for 10min at RT, washed, and kept in PBS + DAPI at 4 °C until imaging on a Nikon A1R confocal microscope. Images were processed and eGFP-positive clusters were counted using Fiji.

### Statistical analysis

Statistical analyses were performed using Graphpad Prism (version 9.0). Data are presented as means ±SEM of 2 or 3 biological replicates (as indicated in the figure legends). One-way ANOVA was used to compare all groups versus control conditions using a Šidák’s multiple comparisons post test.

---

**Supplementary Table 1.**
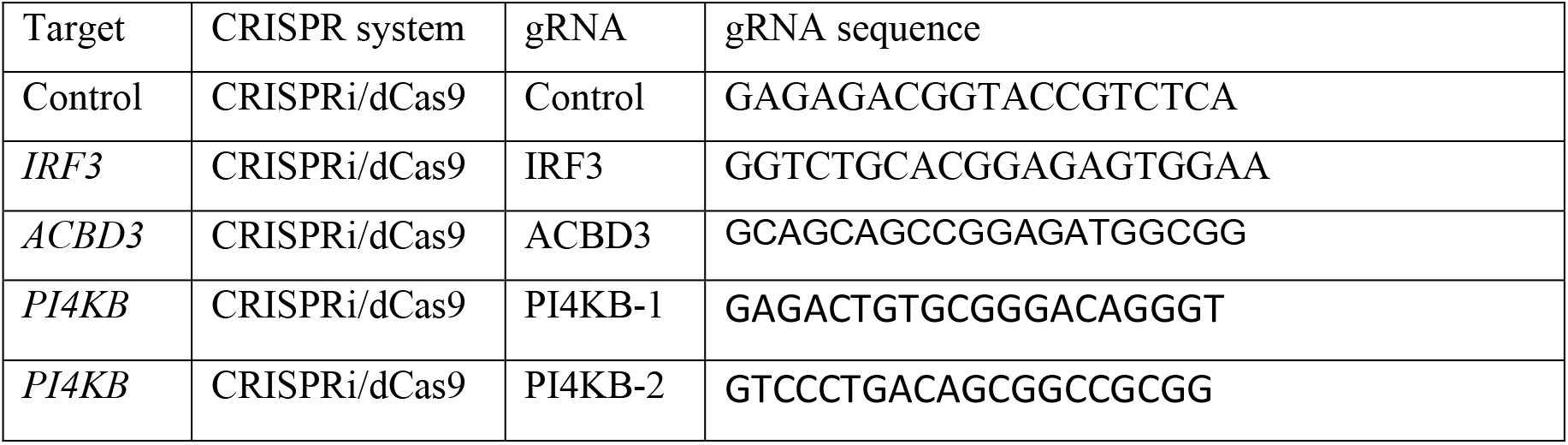
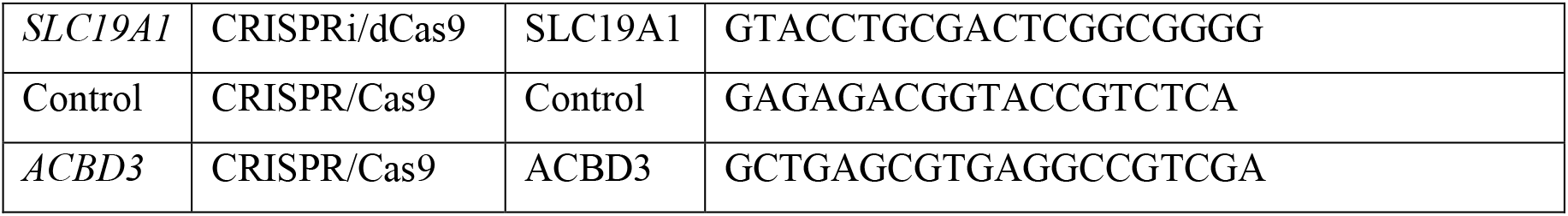
gRNAs used in this study.

**Supplementary Table 2.**
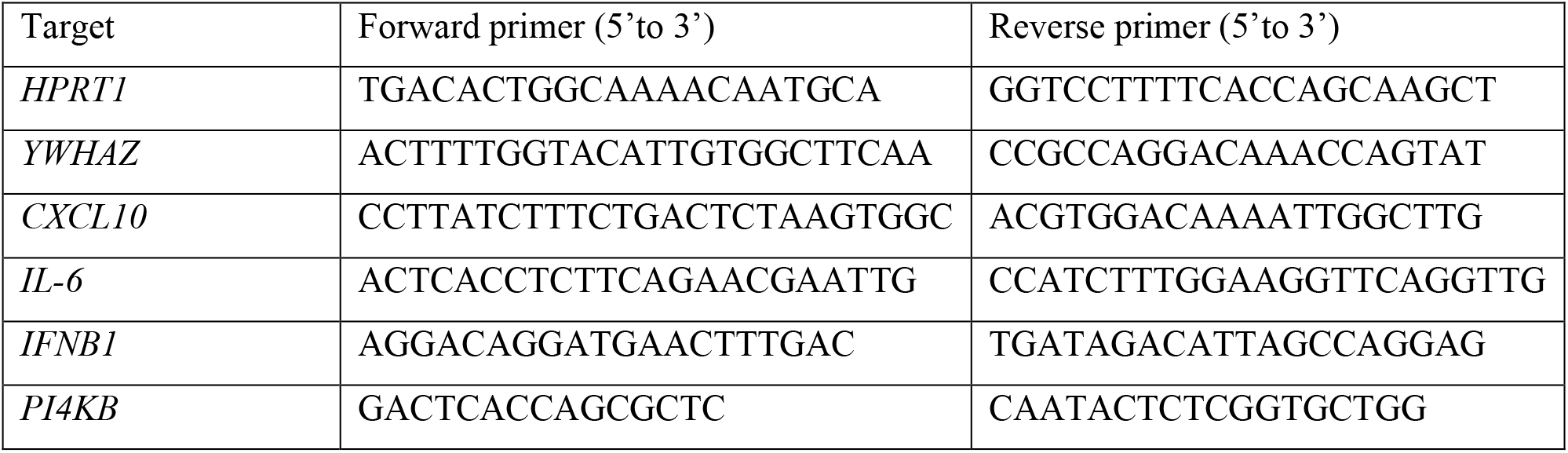
Primers used for RT-qPCR.

Figure S1. ACBD3 expression in THP-1 knockout and knockdown cells.

a. Immunoblot analysis of protein expression in THP-1 cells expressing indicated CRISPRi gRNAs.

b. immunoblot analysis of a THP-1 WT clone and 2 ACBD3 KO clones produced using the conventional CRISPR-Cas9 system

a-b. Representative images of *n* = 2 biological replicates are shown.

Figure S2. ACBD3 knockout in 293 T cells inhibits STING signaling induced by 2’3’-RR CDA

a. tdTomato reporter expression of a 293T WT clone and 2 ACBD3 KO clones stimulated with a. 2’3’-RR CDA (1.7μg/ml) or b. human interferon beta (hIFNb). Reporter activation was measured 20-24h after stimulation. Mean ± SEM of *n* = 3 biological replicates are shown.

Figure S3. After activation, STING is recruited to colocalize with ACBD3.

a. Immunofluorescence images of PMA-differentiated THP-1 cells expressing control gRNA or ACBD3 gRNA. Cells were stimulated for 2h with 5μg/ml 2’3’-RR CDA and stained for ACBD3 and p-STING. Representative images of n = 2 biological replicates are shown.

Figure S4. OSBP inhibitors itraconazole and OSW-1 enhance STING pathway activation.

a. Immublot analysis of indicated (phosphorylated) proteins expressed by THP-1 cells. Cells pre-incubated for 1h with DMSO (D), itrazonazole (I), or OSW-1 (O) were left unstimulated (no stim), or were stimulated with 2’3’-cGAMP (cGAMP; 20μg/ml) for indicated time points

b. immunoblot analysis of THP-1 cells pre-incubated as in panel a and stimulated for 8h with 2’3’-cGAMP (20μg/ml). a-b. Representative images of n = 3 biological replicates are shown.

c. 2’3’-cGAMP uptake (measured by ELISA) in THP-1 cells stimulated or not with 2’3’-cGAMP in the presence of DMSO or itraconazole (ITZ). Mean ±SEM of *n* = 2 biological replicates are shown.

## Notes

### Competing Interest Statement

The authors have declared no competing interest.

