## supplemental files for "STING activation depends on ACBD3 and other phosphatidylinositol 4-phosphate-regulating proteins"

Figure S1. ACBD3 expression in THP-1 knockout and knockdown cells.

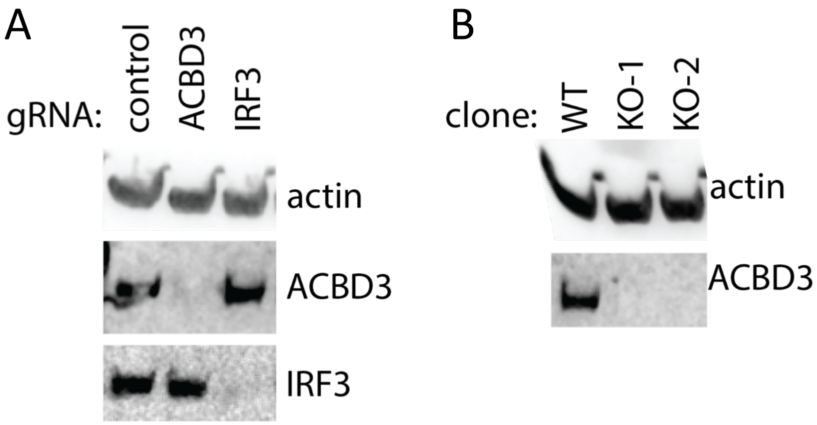


Figure S1. ACBD3 expression in THP-1 knockout and knockdown cells.

a. Immunoblot analysis of protein expression in THP-1 cells expressing indicated CRISPRi gRNAs.

b. immunoblot analysis of a THP-1 WT clone and 2 ACBD3 KO clones produced using the conventional CRISPR-Cas9 system

a-b. Representative images of *n* = 2 biological replicates are shown.

Figure S2. ACBD3 knockout in 293 T cells inhibits STING signaling induced by 2’3’-RR CDA

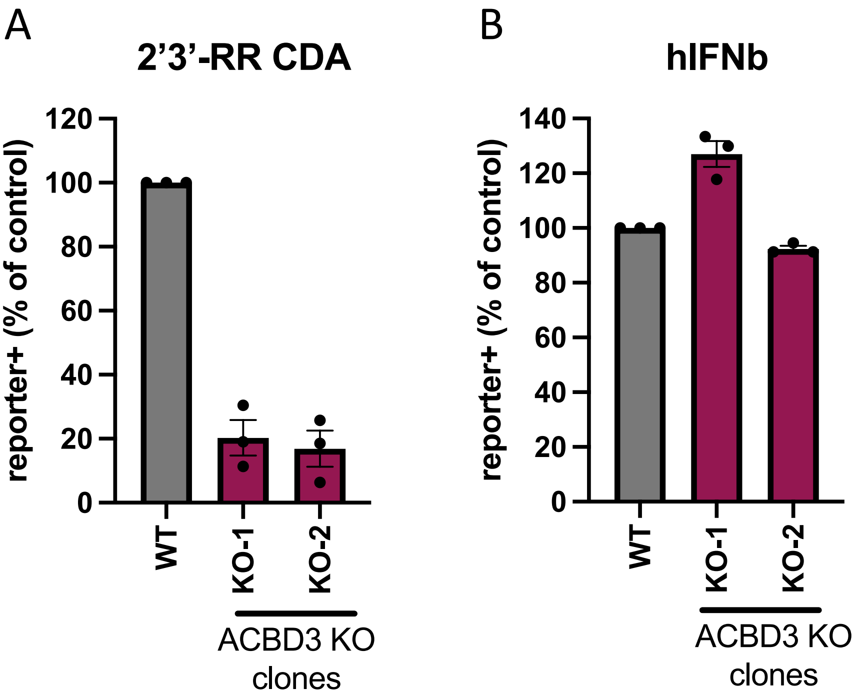


Figure S2. ACBD3 knockout in 293 T cells inhibits STING signaling induced by 2’3’-RR CDA

a. tdTomato reporter expression of a 293T WT clone and 2 ACBD3 KO clones stimulated with a. 2’3’-RR CDA (1.7µg/ml) or b. human interferon beta (hIFNb). Reporter activation was measured 20-24h after stimulation. Mean ± SEM of $n$ = 3 biological replicates are shown.

Figure S3. After activation, STING is recruited to colocalize with ACBD3.

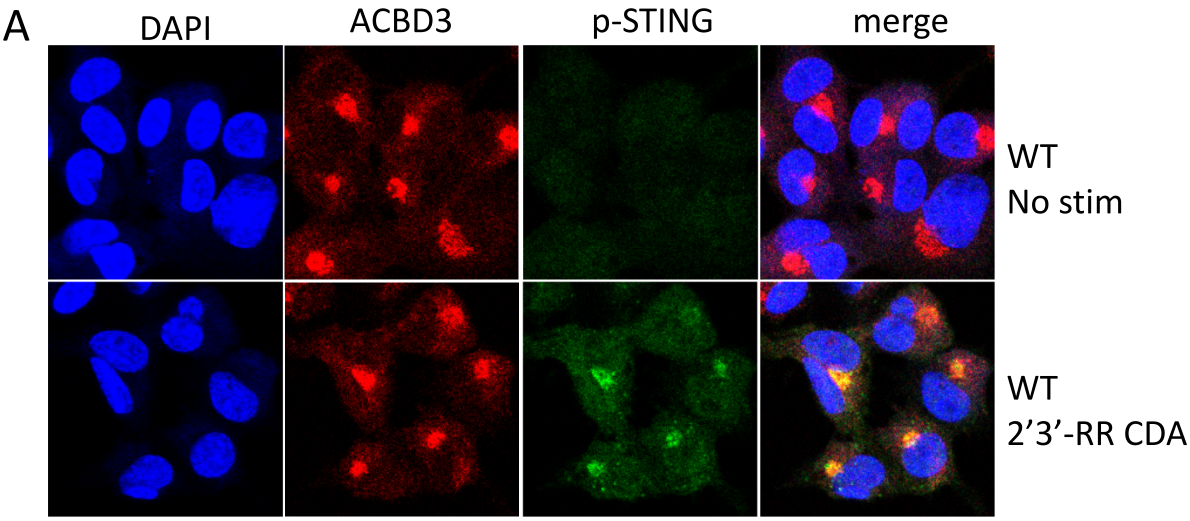


Figure S3. After activation, STING is recruited to colocalize with ACBD3.

a. Immunofluorescence images of PMA-differentiated THP-1 cells expressing control gRNA or ACBD3 gRNA. Cells were stimulated for 2h with 5µg/ml 2'3'-RR CDA and stained for ACBD3 and p-STING. Representative images of n = 2 biological replicates are shown.

Figure S4. OSBP inhibitors itraconazole and OSW-1 enhance STING pathway activation.

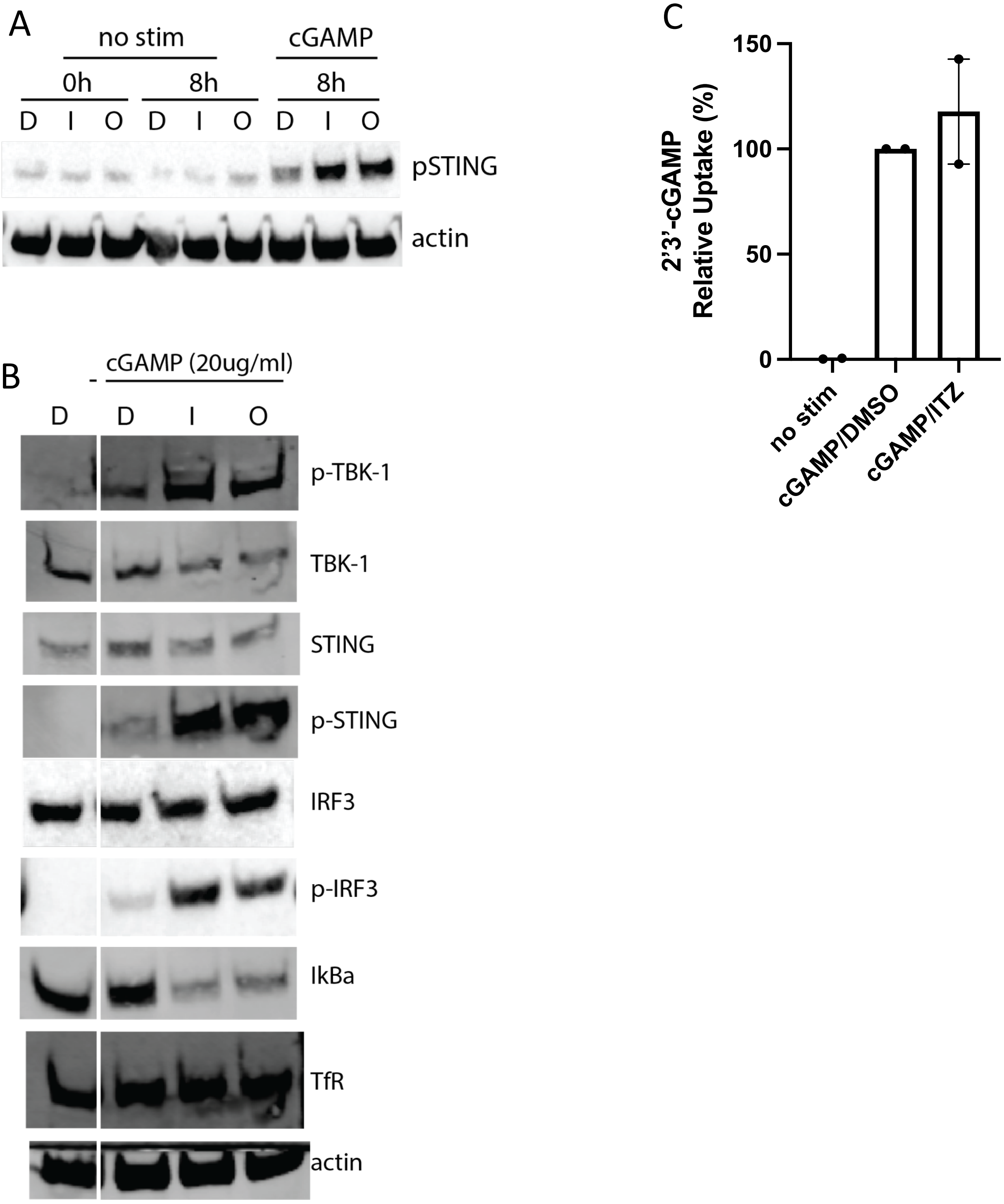


Figure S4. OSBP inhibitors itraconazole and OSW-1 enhance STING pathway activation.

a. Immublot analysis of indicated (phosphorylated) proteins expressed by THP-1 cells. Cells pre-incubated for 1h with DMSO (D), itrazonazole (I), or OSW-1 (O) were left unstimulated (no stim), or were stimulated with 2'3'-cGAMP (cGAMP; 20μg/ml) for indicated time points

b. immunoblot analysis of THP-1 cells pre-incubated as in panel a and stimulated for 8h with 2'3'-cGAMP (20μg/ml).
a-b. Representative images of n = 3 biological replicates are shown.

a. 2'3'-cGAMP uptake (measured by ELISA) in THP-1 cells stimulated or not with 2'3'-cGAMP in the presence of DMSO or itraconazole (ITZ). Mean ±SEM of $n = 2$ biological replicates are shown.
